# UCS: a unified approach to cell segmentation for subcellular spatial transcriptomics

**DOI:** 10.1101/2024.07.08.601384

**Authors:** Yuheng Chen, Xin Xu, Xiaomeng Wan, Jiashun Xiao, Can Yang

## Abstract

Subcellular Spatial Transcriptomics (SST) represents an innovative technology enabling researchers to investigate gene expression at the subcellular level within tissues. To comprehend the spatial architecture of a given tissue, cell segmentation plays a crucial role in attributing the measured transcripts to individual cells. However, existing cell segmentation methods for SST datasets still face challenges in accurately distinguishing cell boundaries due to the varying characteristics of SST technologies. In this study, we propose a unified approach to cell segmentation (UCS) specifically designed for SST data obtained from diverse platforms, including 10X Xenium, NanoString CosMx, MERSCOPE, and Stereo-seq. UCS leverages deep learning techniques to achieve high accuracy in cell segmentation by integrating nuclei segmentation from nuclei staining and transcript data. Compared to current methods, UCS not only provides more precise transcript assignment to individual cells but also offers computational advantages for large-scale SST data analysis. The analysis output of UCS further supports versatile downstream analyses, such as subcellular gene classification and missing cell detection. By employing UCS, researchers gain the ability to characterize gene expression patterns at both the cellular and subcellular levels, leading to a deeper understanding of tissue architecture and function.

## Introduction

Understanding how different cell types assemble into tissues and organs, how cells interact to transmit and receive biological signals, and how cellular communities make decisions in their spatial context are fundamental questions in biomedical and biological research. Subcellular Spatial Transcriptomics (SST) offers a powerful method to measure gene expression at the subcellular level within intact tissue samples, offering unprecedented opportunities to explore the spatial organization and dynamic functions of tissues [1][2][3][4]. To deepen our understanding of cellular behavior using SST technologies, it is essential to accurately attribute the measured transcripts to individual cells through the segmentation of the spatial domain. The precision of cell segmentation directly impacts various downstream tasks in SST data analysis, including gene expression quantification, cell type annotation, and quantitative assessment of the cellular environment. For example, if cell segmentation is oversized, it may include transcripts from neighboring cells or extracellular regions, introducing noise that can distort the spatial gene expression patterns and subcellular gene classification. Conversely, fragmented or underestimated segmentation can result in the loss of significant numbers of transcripts, leading to incomplete or inaccurate gene expression profiles and a subsequent loss of valuable information about cellular behavior. Therefore, reliable and robust cell segmentation is crucial for extracting valuable biological insights from SST datasets.

Currently, SST technologies exhibit notable differences in their capabilities across key parameters such as resolution, capture efficiency, and gene profiling capacity, which are critical considerations for their application in various biological research areas [5]. It poses a significant challenge to develop cell segmentation methods that are applicable to multiple SST platforms. Among these platforms, imaging methods like Vizgen’s MERSCOPE [6] and 10X Genomics’ Xenium [7] achieve high resolutions of approximately 100 nm while maintaining capture rates exceeding 90%. This level of resolution is crucial for applications requiring detailed spatial analysis at the cellular or subcellular level. Alternatively, sequencing-based methods offer distinct capabilities. For example, Stereo-seq [8] enables mRNA collection on an array with a fixed spatial resolution of about 0.5 *µm* and a specific capture efficiency of 12,661 transcripts per 100 *µm*^2^ [9]. The gene profiling capacity also varies among these technologies. Platforms like Xenium and MERSCOPE generally focus on profiling hundreds of genes, making them suitable for targeted studies. In contrast, NanoString’s [10] technology enables the profiling of thousands of genes, providing broader coverage. Meanwhile, Stereo-seq stands out by offering whole transcriptome analysis, capturing over 10,000 genes, which is ideal for applications requiring extensive gene expression profiling. Adaptability to the diverse characteristics of these technologies is crucial in order to achieve accurate and reliable cell segmentation.

Considerable efforts have been dedicated to the development of cell segmentation methods for various SST platforms. To facilitate discussion, we can broadly categorize these methods into two groups: transcript-based methods and image-based methods. Transcript-based methods primarily leverage the abundant transcript information from SST datasets. For example, BIDCell [11] achieves highly accurate segmentation by utilizing extensive datasets that include single-cell RNA sequencing reference data, positive marker genes, and even elusive negative marker genes. This method is particularly effective in accurately delineating boundaries for elongated cell types like fibroblasts. However, the extensive information requirements of BIDCell make it inconvenient to use in practical settings. JSTA [12] employs a novel cell-level classifier that assigns cell (sub)types to pixels, facilitating segmentation. Nonetheless, JSTA is often limited to small tissue sections and heavily relies on single-cell RNA information, limiting its scalability and broad applicability. SCS [13] assigns spots to cells by adaptively learning their position relative to the cell center using a transformer neural network, achieving high accuracy. However, SCS may encounter potentially unstable training phases and exhibit some cross-nuclei segmentation. Baysor [14] offers a unique approach by spatially clustering observed molecules to cells and modeling each cell as an ellipsoid using a Gaussian distribution of transcript composition. Although innovative, Baysor is often sensitive to parameter settings and may segment single cells into multiple smaller cells due to varying gene clusters inside, making its practical use challenging compared to faster deep learning methods. On the other hand, image-based methods rely solely on stain images, particularly nuclei staining, which are often the only staining available in SST datasets. Commercial platforms like 10X and Vizgen employ these methods, but they often face challenges such as inaccurately large cells resulting from nuclei dilation in Xenium’s default segmentation or loss of some transcript data due to smaller segmented cells in Vizgen’s approach. Another example is Cellpose [15], a general method for cell segmentation that primarily utilizes various staining images to delineate cell boundaries. When applied to SST datasets, this approach fails to incorporate SST data and only has nuclei staining as the input, potentially leading to less informative and accurate segmentation outcomes.

In this paper, we introduce a unified approach to cell segmentation (UCS) for SST datasets. The advantages of UCS over existing methods are threefold. First, UCS integrates accurate nuclei segmentation results from nuclei staining with the transcript data to predict precise cell boundaries, thereby significantly improving the segmentation accuracy. The combination of nuclei segmentation and transcript data offers a comprehensive perspective that enhances cell segmentation. Nuclei segmentation provides a precise identification of cell centers, which is critical for accurately locating individual cells within a complex tissue structure. On the other hand, transcript data delineates the spatial distribution of gene expression, which helps in determining the cell boundaries more accurately. Second, UCS is applicable to a wide range of SST platforms, including 10X Xenium, NanoString CosMx, MERSCOPE, and Stereo-seq. Third, UCS stands out for its computational efficiency and user-friendliness, in contrast to methods like BIDCell or JSTA, which require extensive inputs such as single-cell RNA data and specific marker gene information. Through real data examples and downstream analysis, UCS demonstrates its power by enhancing the accuracy of cell segmentation. It produces better visualization results, matching the hematoxylin and eosin (H&E) staining. Moreover, UCS improves downstream analysis by enabling more accurate spatial gene analysis, facilitating subcellular gene classification, and accommodating diverse cell morphologies. The UCS software is now publicly available at: https://github.com/YangLabHKUST/UCS.

## Results

### Method overview

UCS is a deep learning-based approach designed to enhance cell segmentation in diverse SST datasets. By integrating both nuclei segmentation from nuclei staining and transcript data, UCS can distinguish the cell boundaries with high precision. UCS is composed of two strategically designed efficient convolutional neural networks one for predicting the foreground, the region with higher transcript density that likely corresponds to cellular areas and the other for distinguishing the exact cell boundaries. More specifically, UCS comprises two steps in real data applications. Initially, it uses the transcript data to identify the foreground region of candidate cells using the first network according to the density of transcripts (Fig. 1**a**). Subsequently, UCS transforms the nuclei segmentation mask into a softmask. The nuclei segmentation mask is derived from nuclei staining and serves as an initial identification of where cell nuclei are located. Each pixel’s value in the softmask is determined by its distance to the nearby nucleus, representing the probability that the pixel belongs to a particular cell. Then the softmask is combined with the segmented foreground regions to generate cell boundaries for each nucleus utilizing the second network (Fig. 1**b**). Intuitively, the nuclei segmentation mask can serve as an anchor for each cell, while the softmask acts as a probabilistic guidance like a prior, guiding the cell segmentation process using the foreground regions detected from the first step. UCS significantly improved the cell segmentations by associating nuclei with their corresponding cytoplasmic regions, and achieves high consistency with cell segmentations from H&E staining image (Fig. 1**c**), facilitating downstream analyses.

**Figure 1:**
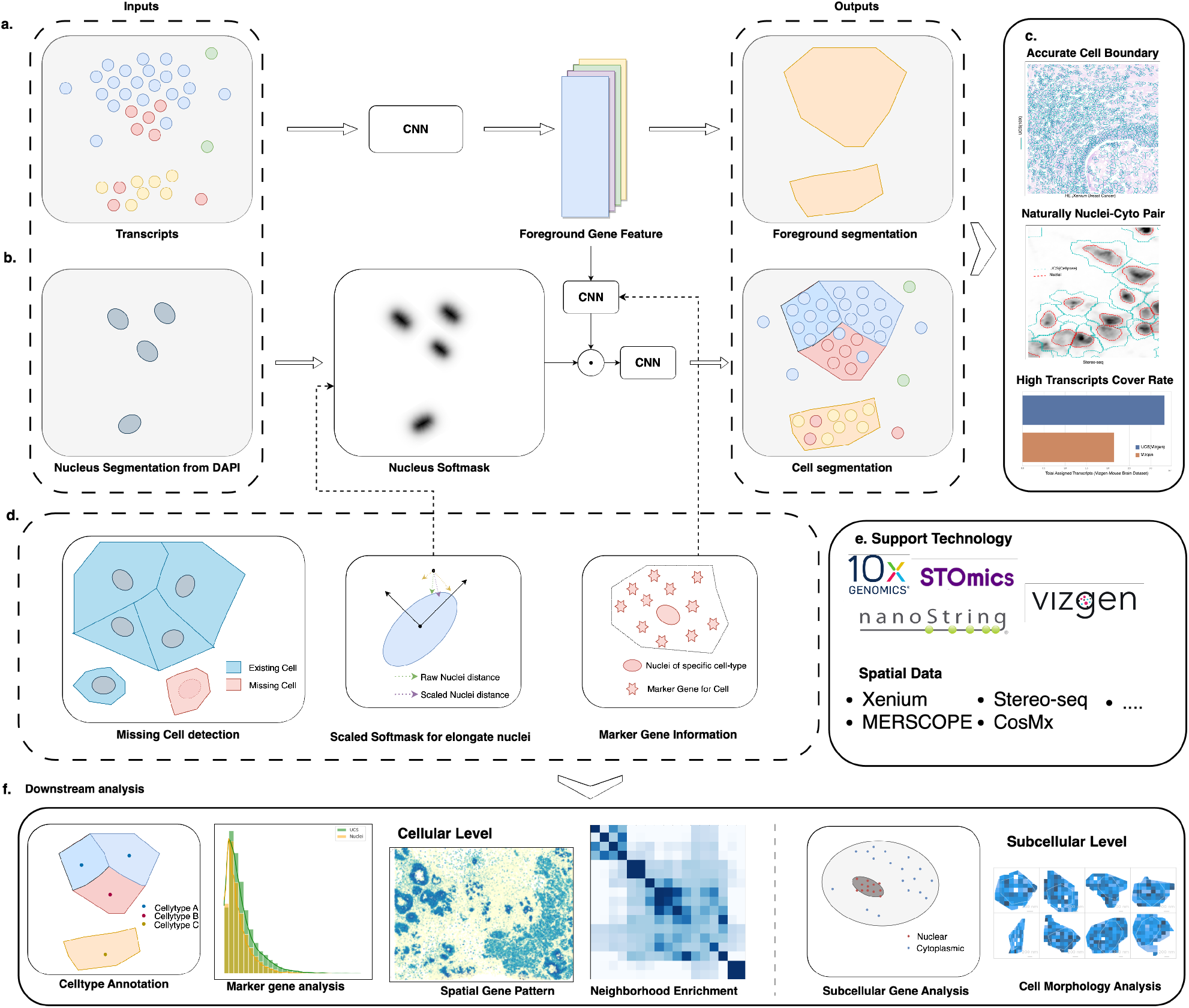
Overview of UCS. **a**. UCS first uses the transcript data as the input and output the potential foreground where cells exist. **b**. Then the nuclei segmentation mask will be processed into a softmask, and integrated with foreground information to get the final cell segmentation mask. **c**. UCS can give accurate cell boundaries and naturally nuclei-cyto pair with a relatively high transcript cover rate. **d**. Other versatile utility of UCS. **e**. UCS can be used on SST datasets from different platforms. **f**. UCS also improves downstream analysis at both cellular level and subcellar level.

UCS is a versatile approach designed to address multiple challenges inherent in cell segmentation task, offering a range of optional utilities that enhance its applicability (Fig. 1**d**). First, UCS exhibits robust detection capabilities, identifying cells that are frequently missed by other segmentation methods. Newly identified cells exhibit sufficient gene expression and compatibility with previously identified cells, ensuring reliable detection. Second, for elongated cell types, UCS employs a scaled softmask to maintain shape consistency with the nuclei, thereby preserving the morphological integrity of these cells. Additionally, UCS can integrate marker gene information to enhance segmentation, ensuring that each nucleus is associated with the correct cell-type specific markers. In summary, UCS is a unified approach that can be applied across various SST datasets generated by different technologies, including 10X Xenium, Vizgen, Nanostring CosMx, and Stereo-seq (Fig. 1**e**). The superior performance of UCS lays the foundation for a range of downstream analyses (Fig. 1**f**). At the cellular level, UCS facilitates cell type annotation, marker gene analysis, spatial pattern recognition, and neighborhood enrichment. At the subcellular level, it supports subcellular gene classification and cell morphology analysis. These capabilities demonstrate the extensive utility of UCS in analyzing SST datasets.

### UCS shows robust performance across diverse SST Datasets

As a general framework that can accommodate various SST datasets through a well-structured training process, UCS can handle the diverse characteristics and varying quality of data across different platforms, ensuring accurate and reliable cell segmentation. Variability within SST datasets, particularly in relation to the tissue type and dataset quality, significantly complicates the process of cell segmentation. For instance, tissues such as breast cancer contain elongated fibroblast cells, which pose unique segmentation challenges compared to other cell types due to their distinct morphology (Fig. 2**a**, I). The quality of SST data also varies across platforms; for example, Vizgen and Nanostring tend to produce cleaner data, whereas Stereo-seq data is often noisier (Fig. 2**a**, II). Additionally, cell density can differ markedly between datasets or even within different regions of the same dataset (Fig. 2**a**, III), necessitating adaptable segmentation methods capable of handling such variability. These challenges underscore the need for robust and versatile cell segmentation techniques that can accommodate the diverse and complex nature of SST data.

**Figure 2:**
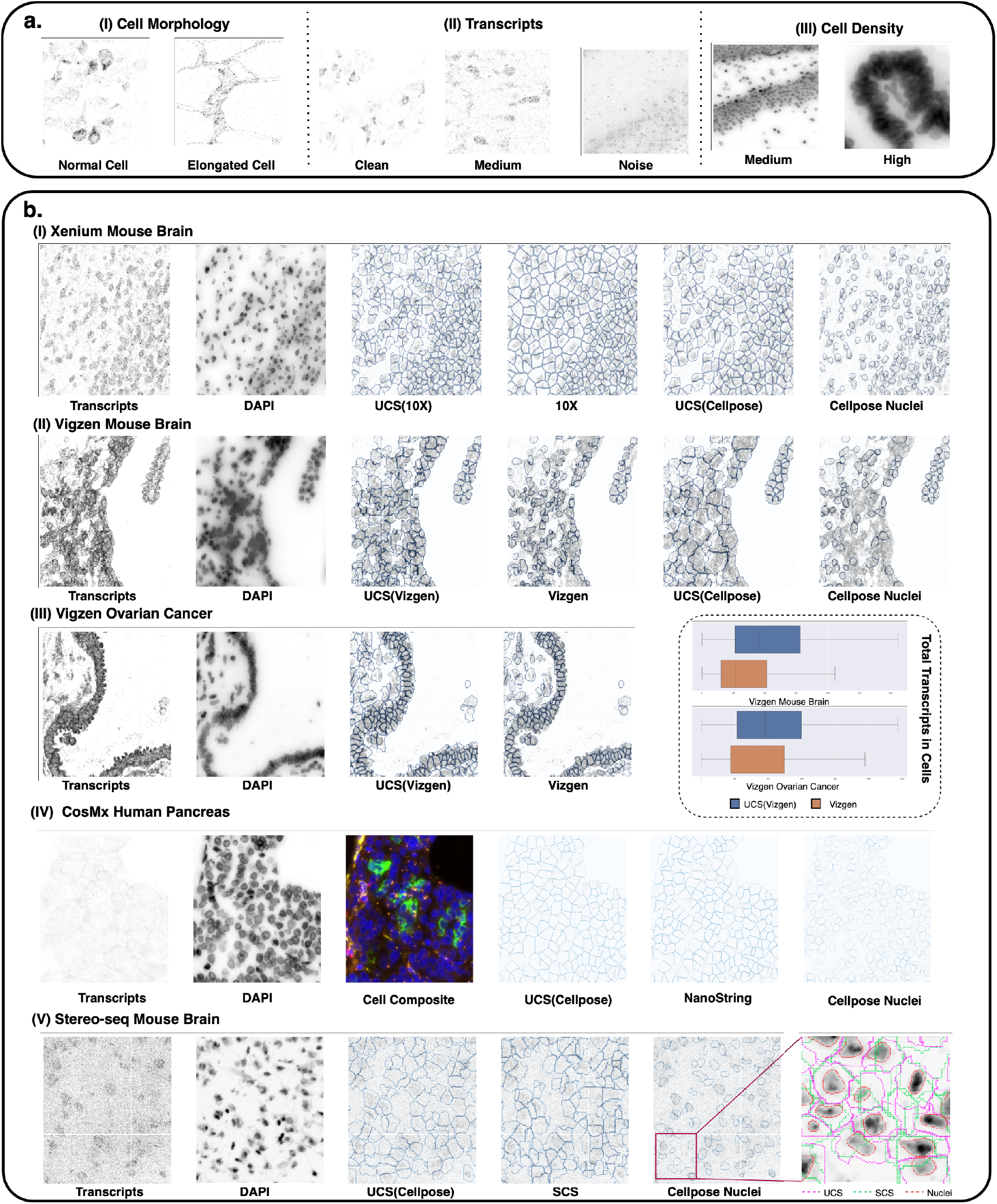
UCS shows robust performance across diverse SST Datasets. UCS (10X) means the nuclei segmentation input for UCS model is produced by 10X, and same notation for UCS (Cellpose) and UCS (Vizgen). **a**. SST datasets vary in cell morphology, transcript noise level and cell density. **b**. Comparative visualization of cell segmentation across diverse SST datasets. UCS shows robust performance across datasets from different platforms. Here, the cell composite image of CosMx Huamn Pancreas dataset is the mixed fluorescence image of nuclei, cytoplasm and cell boundaries.

The proposed UCS method demonstrates superior performance in cell segmentation across diverse SST datasets. Comparative analysis on Xenium Mouse Brain indicates that UCS achieves higher accuracy and produces more compact segmentation results compared to the 10X method (Fig. 2**b**, I), which uses dilation of the nuclei segmentation that often results in oversized cells with low morphological diversity. By leveraging the underestimated segmentation results from Vizgen as “nuclei”, UCS improves transcript coverage rates and includes areas with high gene expression levels (Fig. 2**b**, II, III), thus increasing transcript counts per cell for downstream analyses. Vizgen, on the other hand, adopts overly conservative segmentation strategies, resulting in low transcript coverage rates. In the CosMx Human Pancreas, UCS produces segmentation boundaries comparable to the latest method of NanoString based on the staining of the whole cell (Fig. 2**b**, IV), despite using only nuclei fluorescence information and transcripts. This is important since most SST datasets only provide nuclei staining, but most of the existing image-based cell segmentation methods solely rely on nuclei staining, making it infeasible to precisely segment the cytoplasm of cells based solely on image data. Although recent advancements in platforms like 10X Genomics Xenium 2.0 and NanoString have integrated more sophisticated fluorescence-based imaging techniques (like the newly released CosMx Human Pancreas dataset in the figure), enabling a richer and more detailed capture of both cellular and subcellular structures, the availability of datasets generated from these new technologies remains relatively limited. In the Stereo-seq Mouse Brain, UCS effectively avoids the problematic cross-nuclei segmentation observed with SCS (Fig. 2**b**, V). The stability of UCS is particularly noteworthy in noisy datasets like Stereo-seq, where methods like SCS exhibit unstable training and segmentation results. UCS consistently provides stable and reliable segmentation across all tested datasets, underscoring its robustness and adaptability to various SST scenarios (Supplementary Fig. 2).

By addressing the limitations of both transcripts-based and image-based segmentation methods, UCS offers a unified approach that enhances cell segmentation accuracy and reliability across a variety of SST data from different platforms, tissues, and sections, paving the way for more precise and insightful SST analyses.

### Benchmarking studies show the superior performance of UCS in cell segmentation on Xenium Datasets

To quantitatively compare the cell segmentation performance of UCS and the other methods, we conducted experiments with UCS on various Xenium datasets, including the Xenium Human Breast Cancer, Xenium Mouse Brain, and the recently introduced Xenium 2.0 Human Lung Cancer datasets. UCS excels in both normal and elongated cell regions within the Xenium Human Breast Cancer dataset (Fig. 3**a**). By leveraging the scaled softmask, marker gene information, and nuclei segmentation provided by Cellpose, UCS achieves segmentation results comparable to BIDCell—a method specifically designed for elongated cell types, which utilizes prior knowledge about the elongation of cells. In comparison, Baysor, which depends on gene clustering, often yields small or fragmented cells. This fragmentation is a consequence of clustering-based approaches that may not fully consider the spatial continuity and morphological context of cells. On the other hand, BIDCell employs extensive additional information to assist in cell segmentation, such as single-cell RNA-seq data and other contextual markers. This limits BIDCell’s applicability. Furthermore, UCS exhibits greater consistency with H&E staining (Fig. 3**b**). The regions with deep color staining, which indicate the presence of cells, are more accurately covered by UCS segmentation than by BIDCell, while blank regions are not erroneously segmented a problem seen in the segmentation results of 10X. UCS also ensures that the boundaries of cell clusters are distinctly clear and demonstrates superior accuracy in cell annotation.

**Figure 3:**
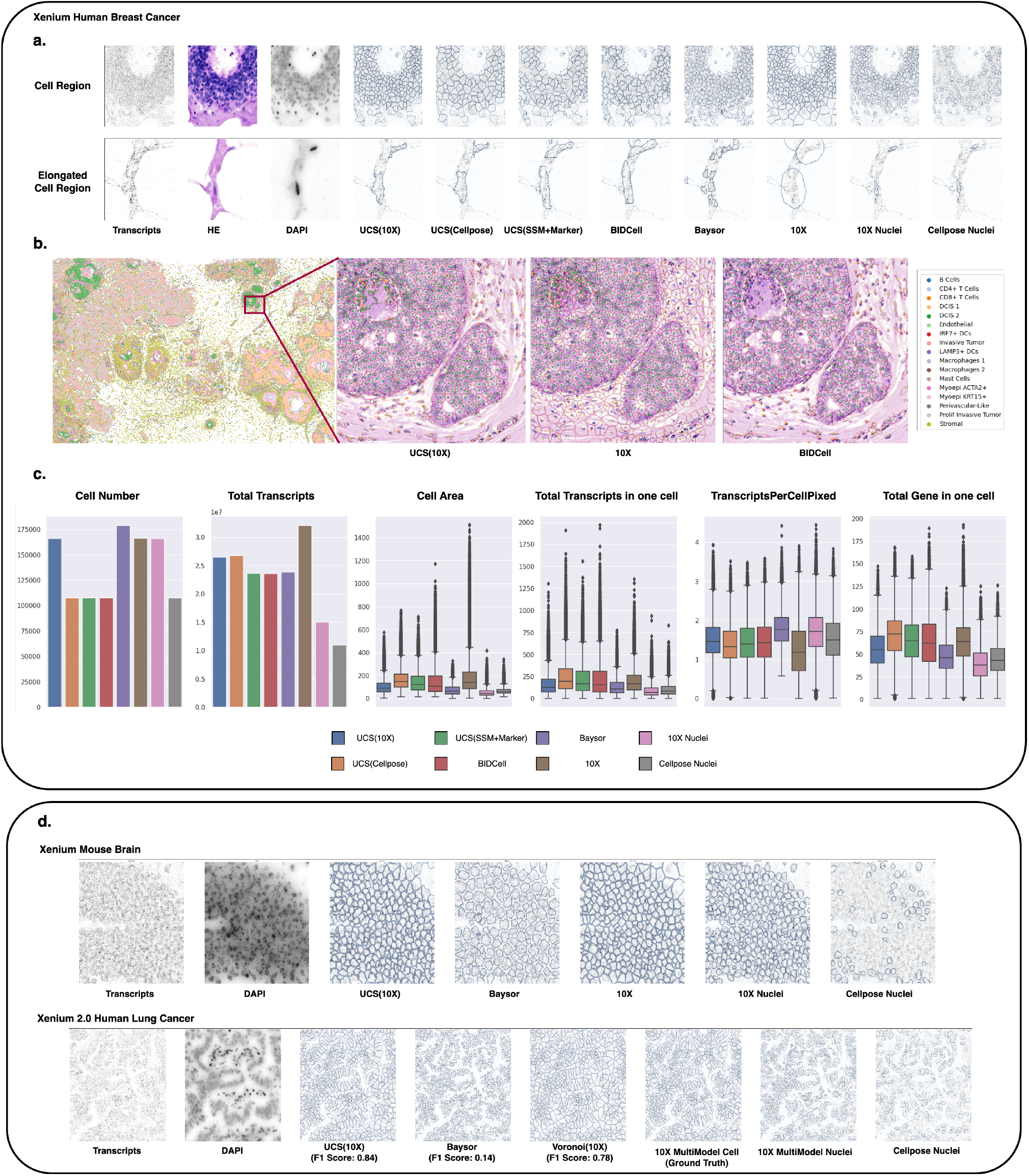
Benchmark using Xenium datasets. UCS(SSM+Marker) means UCS using scaled softmask, additional marker gene information(discussed in detail in the Method section) and Cellpose nuclei segmentation as the input. **a**. Comparison of normal cell region and elongated cell region with different methods on Xenium Human Breast Cancer dataset. The H&E images are provided for visual comparison. **b**. Cell annotation result on Xenium Human Breast Cancer dataset. UCS shows greater consistency with the H&E image which is not used for building the segmentation model compared to 10X or BIDCell. **c**. Cell segmentation characteristics of different methods. **d**. More comparisons on Xenium Mouse Brain dataset and Xenium 2.0 Human Lung Cancer dataset. UCS has the largest F1 Score with the 10X MultiModel Cell which uses additional fluorescence information.

To establish baseline characteristics, we computed several metrics across different methods, including cell number, total transcripts, cell area, total transcripts in one cell, transcripts per cell pixel, and total genes in one cell (Fig. 3**c**). It is important to note that these metrics alone do not guarantee an unbiased comparison, as larger cells naturally encompass more transcripts, and vice versa. The analysis reveals that BIDCell tends to produce more large “outlier” cells, while 10X methods result in larger cells without a corresponding increase in transcript count, suggesting potential oversizing in segmentation, which is consistent with previous observation in Fig. 3**b**.

In addition to the Human Breast Cancer dataset, we further evaluated UCS on the Xenium Mouse Brain and the new Xenium 2.0 Human Lung Cancer datasets (Fig. 3**d**). The Xenium 2.0 dataset employs advanced fluorescence technology to stain both nuclei and cell boundaries, providing a more convincing ground truth for cell segmentation evaluation. Using the 10X Multimodel result for benchmarking, we calculated the F1 Score for various methods (Fig. 3**d**) [16]. UCS achieved the highest F1 Score of 0.84, indicating its segmentation results are closely aligned with the ground truth provided by the 10X 2.0 dataset.

### UCS Enhances the Consistency of Cellular Expression Profiles with scRNA-seq Data

An accurate cell segmentation method will generate cellular gene expression profiles that closely match the corresponding scRNA-seq reference dataset. Given this fact, we integrated the spatial single cells segmented by the compared methods with a scRNA-seq dataset from the same tissue. We evaluated the integration result with two aspects. From the cell type labeling aspect, we transferred the cell type labels from scRNA-seq to the segmented cells by different methods and then assessed their spatial organizations. As shown in Fig. 4**a**, UCS enabled a more precise identification of the spatial distribution of cell types in human breast cancer tissue, especially accurately distinguishing the two subtypes (Fig. 4**a**) (Green and Light orange regions) in areas of ductal carcinoma in situ (DCIS). While different segmentation methods generally reflected similar proportions of each cell type, the 10X segmentation approach often resulted in oversized cell boundaries. This oversizing tends to incorporate more extraneous genetic noise, thereby affecting cell type annotations.

**Figure 4:**
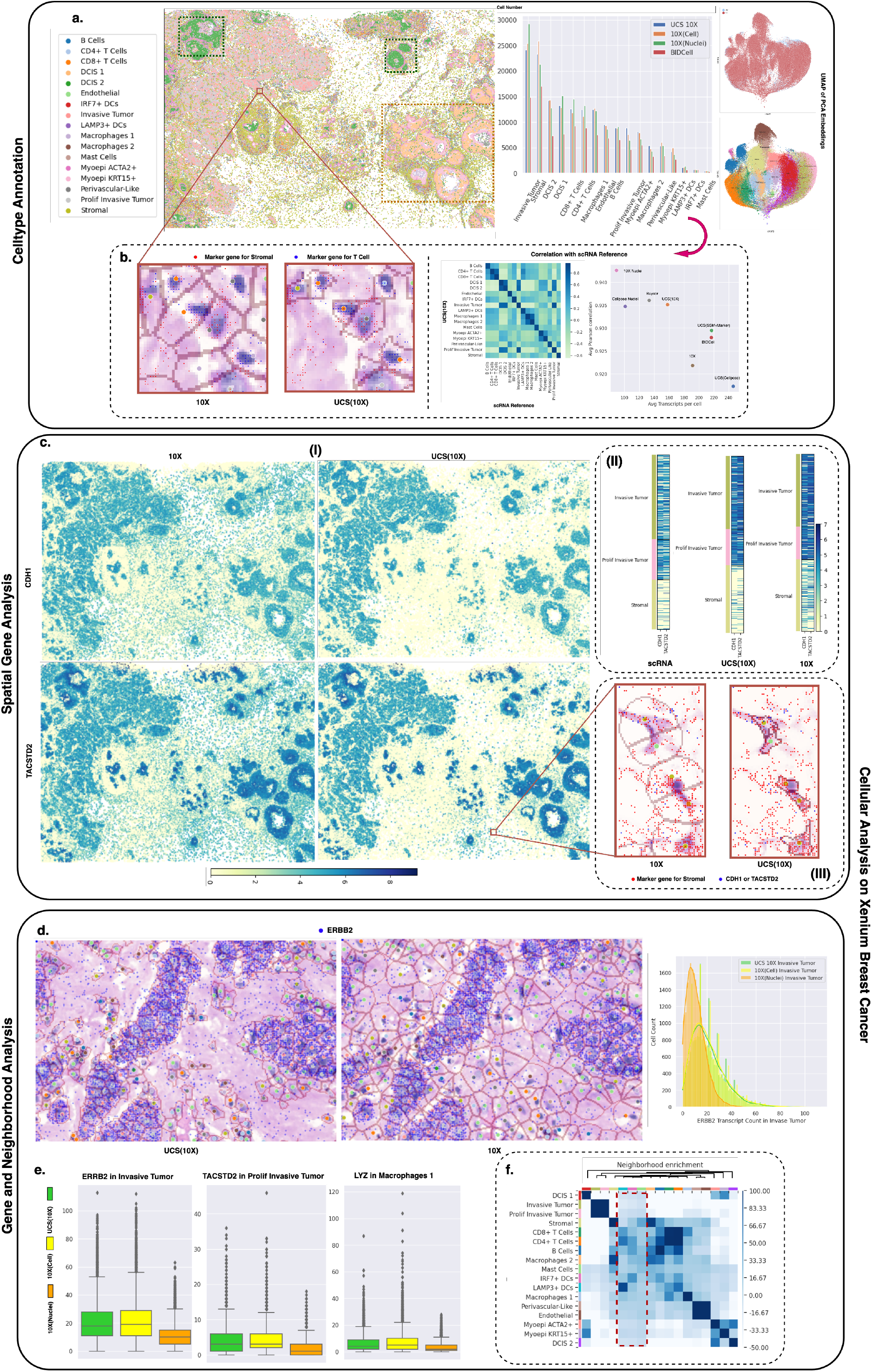
Cellular level analysis. **a**. Cell type annotation results of UCS for Xenium Human Breast Cancer, alongside a joint UMAP visualization of PCA embeddings of both the scRNA dataset and the spatial dataset, demonstrate well-integrated cell expression profiles from the UCS segmentation and the paired scRNA dataset. **b**. Left: A region demonstrates the superior alignment of UCS segmentation with H&E images compared to 10X. UCS includes fewer irrelevant genes, resulting in more accurate cell type annotation. Right: Correlation with scRNA reference. UCS with 10X nuclei achieves high average correlation, maintaining relatively large cell sizes. **c**. The spatial patterns of CDH1 and TACSTD2, predominantly expressed in tumor cells, are clearer and cleaner with UCS compared to 10X. This improvement is due to UCS excluding noise genes that are distant from cell nuclei and located in sparse transcript regions. UCS also shows better consistency in cell expression with the scRNA dataset. **d**. UCS identifies nearly the same highly expressed genes or marker genes in different cells, albeit with relatively smaller but more refined cell sizes compared to 10X. Additionally, noticeable spatial neighborhood enrichment patterns are evident in UCS’s segmentation results.

For gene expression aspect, we compared the Pearson correlation (PCC) between the gene expression of segmented cells of each cell type and corresponding cells from the scRNA-seq reference. As shown in right panel of Fig. 4**b** and Fig. 4**c** II, the cell expression profiles obtained using UCS demonstrated an improvement in similarity with scRNA-seq reference by maintaining appropriate cell sizes and accurate cell boundaries. The enhanced segmentation accuracy of UCS resulted in gene expression profiles that more closely matched those observed in scRNA-seq datasets, yielding cleaner and clearer spatial gene patterns such as *CHD1* and *TACSTD2* (Fig. 4**c** I), which are highly expressed in tumor cells. Additionally, UCS enabled the identification of highly expressed or marker genes in cells, such as *ERBB2* in invasive tumors and *LYZ* in macrophages (Fig. 4**e**). The precise boundary delineation ensured that the number of genes identified was almost the same as with the oversized 10X segmentation results (Fig. 4**d**).

Furthermore, UCS’s segmentation results facilitated the discovery of neighborhood enrichment patterns in breast cancer tissues (Fig. 4**f**). For instance, our analysis revealed that tumor cells inhibit the growth of neighboring cells and suggested potential interactions between immune cells and IRF7+ DCs. These findings underscore UCS’s utility in elucidating complex cellular interactions within the tissue microenvironment, thereby advancing our understanding of spatially resolved transcriptomics.

### UCS Enhances Subcellular Gene Localization and Morphological Correlation

To further illustrate the utility and advantage of UCS in subcellular gene classification, we employed Bento [17] to determine the primary expression locations of genes—nuclei, cytoplasm, or no preference. We specifically analyzed an invasive tumor region with a cell count comparable to the dataset used in the original Bento paper (Fig. 5**a**, left). Notably, in both UCS and 10X segmentation results, the top three genes expressed in the nuclei were *ANKRD30A, POLR2J3*, and *MLPH*. However, discrepancies emerged in the cytoplasmic gene classification: UCS identified *SMS, HMGA1*, and *SEC11C*, whereas 10X identified *LLM, SEC11C*, and *FAM107B*, with *LLM* showing significantly higher expression levels (Fig. 5**a**, right). Upon further investigation using scRNA-seq references, *LLM* was found to be highly expressed in stromal cells but rarely in tumor cells, indicating a misclassification by the 10X method due to incorrect cell boundary assignments. In contrast, *SMS*, which UCS identified, is moderately expressed in tumor cells and predominantly localized to the cytoplasm, corroborating the accuracy of UCS segmentation.

**Figure 5:**
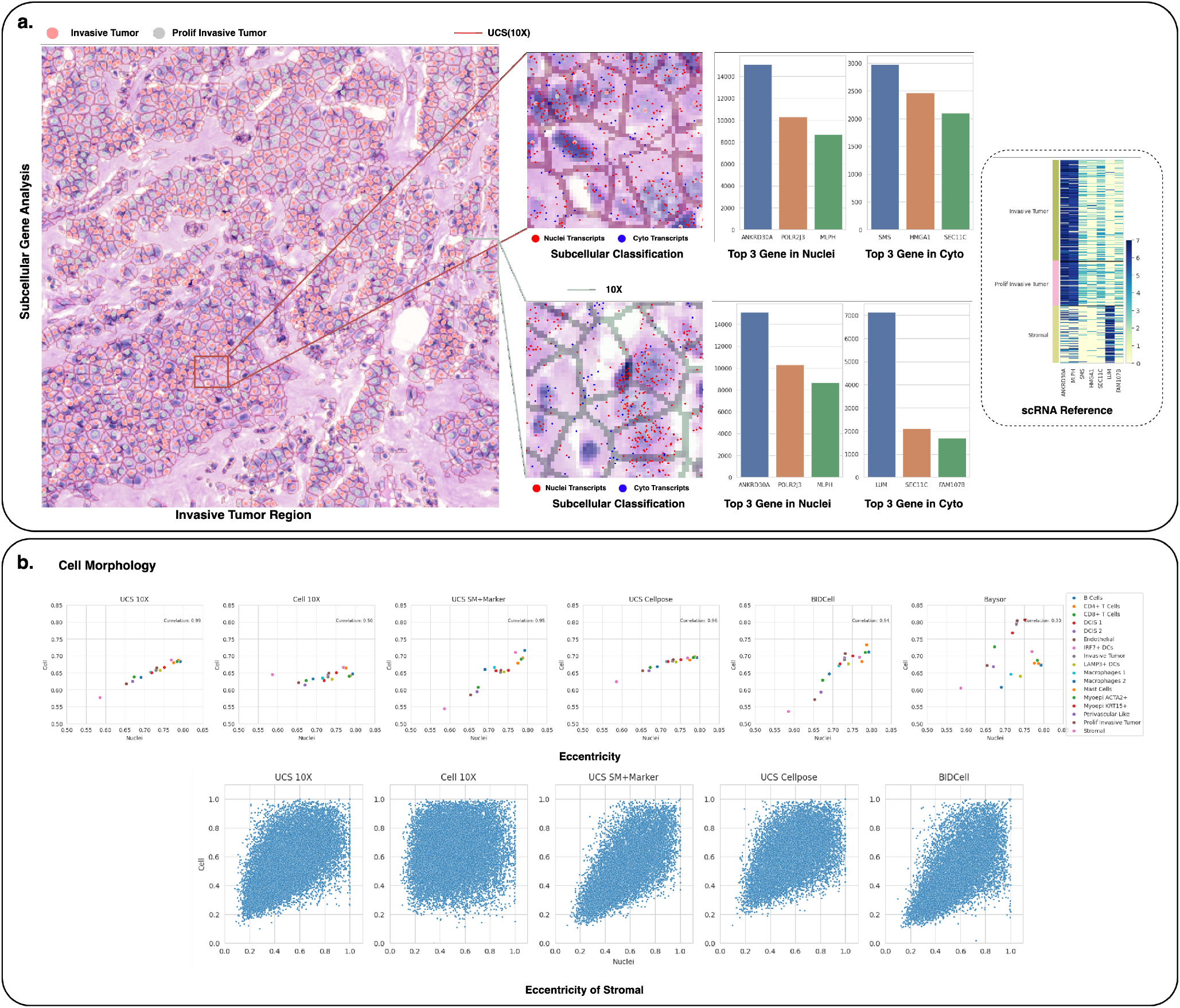
Subcellular level analysis. **a**. Subcellular classification results in an invasive tumor region. Genes are classified based on their primary expression location—nuclei, cytoplasm, or no preference—and the top three genes expressed in the cytoplasm and nuclei are listed for both UCS and 10X segmentation results. UCS correctly identifies the relevant genes, whereas 10X misclassifies genes from other cell types as cytoplasmic genes of the tumor, primarily due to its inaccurate boundary definitions. **b**. Eccentricity of the nuclei (x-axis) versus cells (y-axis) to compare cell morphology across methods for each cell type, highlighting the elongated morphology of stromal cells.

Moreover, we compared cell morphology by calculating the eccentricity for nuclei and cells across different cell types (Fig. 5**b**, upper). Both UCS and BIDCell demonstrate the ability to preserve diverse and accurate cell morphology, whereas other methods failed. A concrete example of stromal cells with notably elongated morphology is shown in the bottom panel of Fig. 5**b**, where UCS successfully segmented these elongated cells with the aid of scaled soft masks and marker gene information, matching the performance of BIDCell, which requires extensive prior knowledge about cell elongation and other marker gene information.

### UCS enables Detection of Missing Cells in SST datasets

One of the significant advancements introduced by UCS is its ability to identify missing cells that traditional image-based segmentation methods often overlook. This capability stems from UCS’s two-stage strategy of cell segmentation, which leverages both transcriptomic and imaging data. In the first stage, UCS predicts the confidence foreground region for cells solely based on the SST data. This transcript map provides a detailed view of gene expression across the tissue sample, independent of the staining intensity or quality. By focusing on the gene expression data, UCS can identify potential cell regions that might not be apparent in imaging data alone. In the second stage, UCS refines the initial predictions using prior segmentation results. Specifically, the unassigned potential cell regions are defined as the difference between the confidence foreground and the final cell segmentation result (Fig. 6**a**). This difference highlights areas where traditional image-based methods, which often rely heavily on nuclei staining, might fail due to extremely dense nuclei fluorescence, as illustrated by the red region in the Vizgen Mouse Brain dataset (Fig. 6**b**).

**Figure 6:**
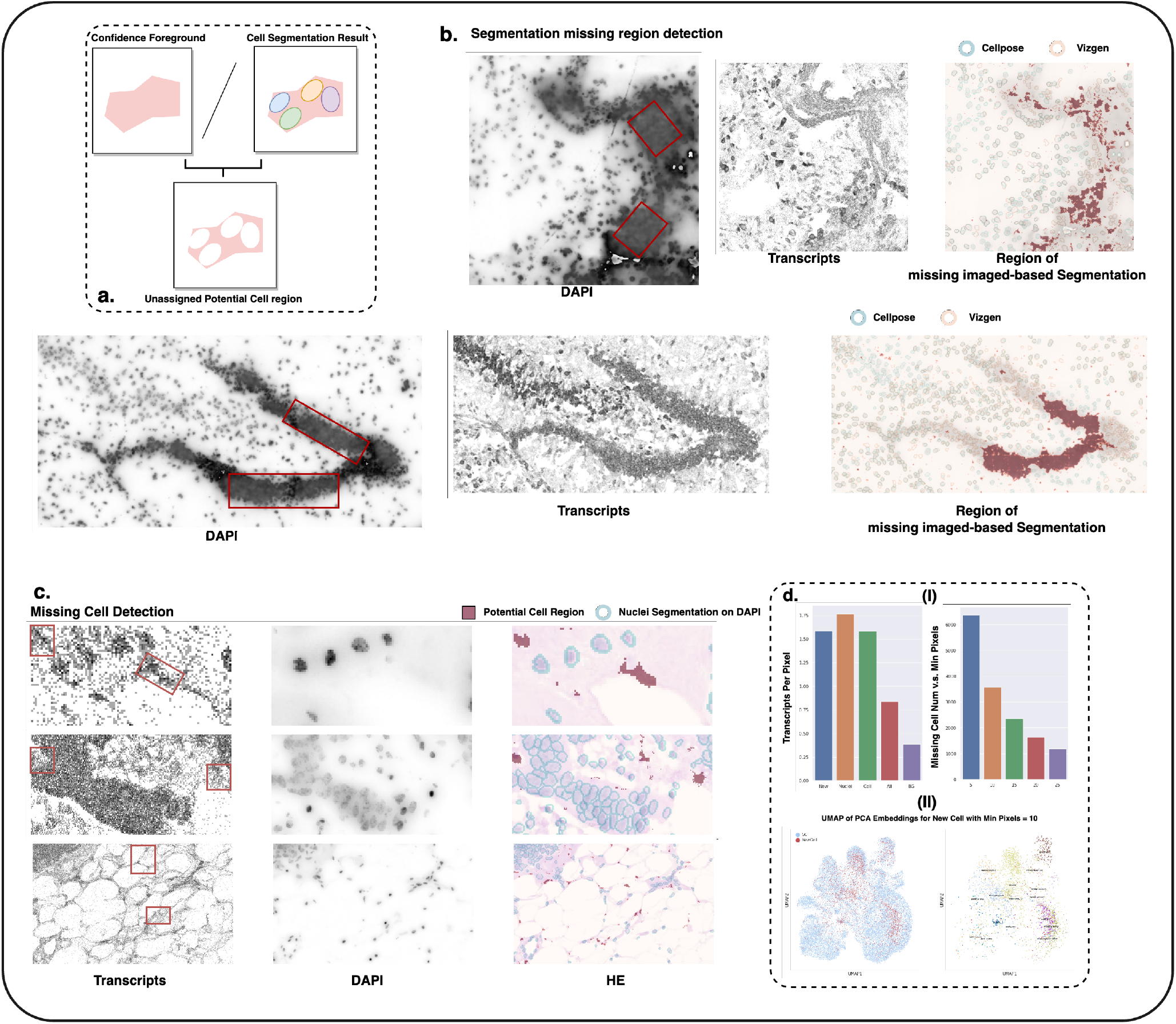
Missing Cell detection. **a**. The unassigned potential cell region is defined as the difference between the confidence foreground and the final cell segmentation result by UCS. **b**. UCS can identify regions where image-based segmentation methods fail, likely due to the extremely dense distribution of different nuclei fluorescence. **c**. Missing cell detection on Xenium Breast Cancer dataset. **d**. UCS can detect missing cells with high gene expression, and the joint UMAP of PCA embeddings of both the scRNA dataset and the detected cells shows that these novel cells integrate well with cells in the scRNA dataset, demonstrating their correctness.

Next, we applied UCS to detect missing cells in less dense spatial data of Xenium Breast Cancer (Fig. 6**c**). We observed that the de nova cells are characterized by transcript counts per pixel that are smaller than those of existing nuclei but larger than those of fully segmented cells (Fig. 6**d**, I). This nuanced detection captures cells that might be underrepresented or entirely missed due to low staining signal intensity in traditional methods. The accuracy of UCS in detecting these novel cells is further validated by integrating their expression profiles into the PCA embedding space (Fig. 6**d**, II). These novel cells’ expression profiles align well with those in corresponding scRNA datasets, indicating the correctness of the UCS-based detections. This integration ensures that the newly identified cells are not artifacts but genuine parts of the cellular landscape.

### Computational time

UCS demonstrates its advantages over existing methods in terms of computational efficiency, making it suitable for large-scale SST data analysis. For the Xenium Breast Cancer dataset, UCS completes the segmentation in approximately 1.5 hours when utilizing nuclei segmentation obtained by Cellpose, processing 107,848 cells. When using 10X nuclei segmentation, the computational time increases slightly to 2 hours for 165,752 cells. Additionally, UCS requires only 12 minutes to segment a 1,200*×*1,200 patch of Stereo-seq and 15 minutes of one 1400*×*1400 Field of view (FOV) of the CosMx Human Pancreas dataset. In contrast, BIDCell, another prominent method, takes around 2 hours to process the Xenium Breast Cancer dataset containing 107,829 cells. Baysor demonstrates a considerably longer computational time. For the Xenium Breast Cancer dataset, Baysor requires approximately 48 hours to complete the segmentation process. The SCS method, when applied to a 1,200*×*1,200 patch of Stereo-seq data, takes about 14 hours to complete the segmentation. All the experiments were conducted on a Single NVIDIA Tesla V100 GPU with a 44-core CPU.

## Discussion

We introduce UCS, a deep learning-based approach designed to enhance cell segmentation within diverse SST datasets. UCS integrates nuclei segmentation from nuclei staining with transcriptomic data, distinguishing cell boundaries with high precision. The method comprises two convolutional neural networks: the first network predicts the foreground regions of candidate cells, while the second network delineates the exact cell boundaries. This two-step process ensures that UCS achieves superior segmentation accuracy, effectively associating nuclei with their corresponding cytoplasmic regions.

The results of employing UCS demonstrate significant advancements in various aspects of SST data analysis. The cell segmentation results of UCS exhibit high accuracy in visual observations and show remarkable consistency with H&E staining images. One of the salient advantages of UCS is its ability to support a broad range of downstream analyses. For example,

UCS substantially enhances the precision of cell type annotation, facilitating more accurate identification and classification of cell types within tissue samples. Moreover, UCS significantly improves the fidelity of gene expression profiles by maintaining appropriate cell sizes and accurate boundaries. This enhancement results in gene expression profiles that closely mirror those obtained from scRNA-seq references, thereby enabling more precise spatial gene pattern recognition and more reliable subcellular gene classification.

Additionally, the improved segmentation accuracy afforded by UCS facilitates the identification of highly expressed or marker genes in specific cell types. UCS’s ability to maintain cell morphology is particularly noteworthy for elongated cell types, such as fibroblasts, which are often challenging to segment accurately with other methods. By preserving the morphological information of these cells, UCS ensures that the resultant gene expression data is both reliable and biologically meaningful.

In conclusion, UCS signifies an advancement in the field of spatial transcriptomics by offering a robust and efficient solution for precise cell segmentation. Its well integration of multimodal data, coupled with its computational efficiency and applicability across a variety of SST platforms, positions UCS as a versatile and powerful tool for researchers. The enhanced segmentation accuracy and downstream analytical capabilities provided by UCS promise to yield more accurate and reliable biological insights, thereby contributing to a deeper understanding of tissue structure and function.

## Methods

### Preprocess SST dataset from different platform

To preprocess SST datasets using the UCS model, we start by converting all detected transcripts with spatial locations into a transcript map. This conversion results in a 3D array denoted as **G** ∈ ℝ^*N×H×W*^, where *N* represents the number of genes in the dataset and *H, W* is the height and the width of transcript map. Each channel in this array corresponds to a specific gene, with each pixel within a channel reflecting the transcript count of that gene at a given location.

For nuclei segmentation, we employ two distinct approaches based on the availability of data from the platform. One approach utilizes the nuclei segmentation data provided directly by platforms such as 10X Genomics Xenium or Vizgen MERSCOPE. These platforms offer pre-segmented nuclei data which can be seamlessly integrated into our preprocessing workflow.

The alternative approach involves using the Cellpose [15] algorithm to segment the nuclei staining. Cellpose is a versatile tool that can handle diverse staining conditions and image qualities, making it ideal for datasets from different platforms.

Following the segmentation, we adjust the segmentation mask to ensure uniform spatial resolution with the transcript map. This step involves resizing the segmentation mask to 0.5 *µm* for datasets from the Stereo-seq and CosMx platforms and to 1 *µm* for datasets from 10X Genomics Xenium and Vizgen MERSCOPE. This resizing ensures the segmentation mask pairs accurately with the transcriptomic data, maintaining the integrity of the size **M** ∈ ℝ^*H×W*^. The final input to the UCS model includes both the transcript map **G** and the resized segmentation mask **M**. By integrating these elements, UCS leverages the combined strengths of transcriptomic data and pre-existing segmentation information. This dual-input strategy enhances the accuracy and robustness of cell segmentation, effectively delineating cell boundaries and identifying individual cells within the tissue.

### The model of UCS

UCS is a deep learning-based method designed for the precise segmentation of cells in diverse spatially resolved transcriptomics data. It comprises two primary neural networks: the Foreground Predict Net and the Cell Predict Net (Supplementary Fig. 6).

### Foreground Predict Net

The Foreground Predict Net focuses on identifying regions within the transcriptomic data that likely correspond to cellular areas. To achieve this, we first divide the whole transcripts map **G** into smaller patches *{***G**^(*p*)^*}* with size *h × w* to facilitate processing (default patch size is 48×48). The input to this network is the transcripts map of a patch **G**^(*p*)^ ∈ ℝ^*N×h×w*^. The Foreground Predict Net consists of:

- A feature extractor *F* (·) with three 1*×*1 kernel convolutional layers.
- An encoder with four 3*×*3 kernel convolutional layers *{e*_*i*_*}, i* = 1, 2, 3, 4. The output of each layer is concatenated to provide a rich and comprehensive feature set. The concatenated output of the encoder is denoted by *E*(·).
- An classifier *C*(·) with four 1*×*1 kernel convolutional layers.

By employing these convolutional layers without any pooling [18], the network ensures that critical gene expression information is not lost, resulting in a robust prediction. Both stride and padding are adjusted to maintain the original dimensions of the patches, thereby preserving the full spatial context of the gene expression data. We can derive the prediction as follows:

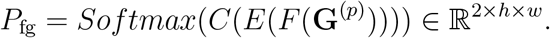

This output has two channels: the first channel represents the probability that a given pixel is part of the background, and the second channel represents the probability that a pixel is part of any cell. We denote 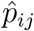 as the predicted probability of the pixel (*i, j*) being foreground.

### Nucleus Softmask Generation

An integral aspect of our UCS model is the generation and utilization of a probability softmask derived from the nuclei segmentation mask. This softmask serves as a crucial input to the Cell Predict Net. To begin, the nuclei segmentation mask *{***M***}* is preprocessed into smaller patches *{***M**^(*p*)^*}*, akin to the division of the transcript map. The nuclei mask for each cell is then expanded into the channel dimension, resulting in an input tensor 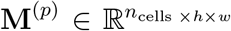. Here, each channel represents a binary mask corresponding to an individual cell’s nuclei.

For each cell’s nuclei segmentation mask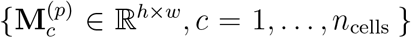, we model the weight of each pixel (*i, j*) using a sigmoid function *σ*(·), which accounts for the pixel’s association with the nuclei [19]. The formula for this transformation is given by:

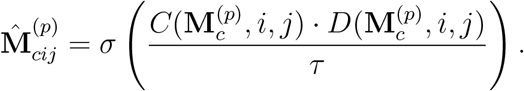

In this equation, 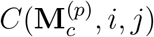 is a value that equals 1 if the pixel (*i, j*) lies within the nuclei of cell *c* and -1 otherwise. 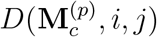 denotes the distance from the pixel to the closest point on the nuclei segmentation boundary. The parameter *τ*, controls the sharpness of the sigmoid function. The resulting softmask 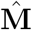 provides a probability distribution over the spatial domain, which is then fed into the Cell Predict Net. The softmask effectively bridges the SST data with the nuclei segmentation information.

### Cell Predict Net

The Cell Predict Net, inspired by the FiLM framework [20], is designed to seamlessly integrate the foreground features derived from the Foreground Predict Net with the softmask to achieve precise cell segmentation. The input to this network is the feature from the Foreground Predict Net

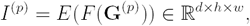

and the softmask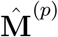. The Cell Predict Net consists of:

- A network *T* (·) with three 1*×*1 kernel convolutional layers designed to process the foreground feature.
- A single 1*×*1 kernel convolutional layer *H*(·) as the predicted head.

The final output for a particular cell *c* in the patch will be:

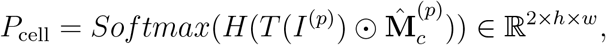

where ⊙ is the element-wise product, effectively weighting the feature map according to the probabilistic information provided by the softmask. This operation ensures that the network focuses on regions with higher probabilities of being part of a cell. The network generates a probability distribution over a cell and background for each pixel. Note that in practice the first dimension (*n*_cells_) of 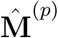 is placed in the batch size dimension, and cells from the same patch use the same foreground feature. We denote 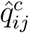 as the predicted probability of the pixel (*i, j*) being cell *c* in the corresponding patch.

To make the final prediction for each patch, consider that there are *C* nuclei of cells within the patch. Given that the batch size is equal to the number of cells, the outputs of the Cell Predict Net for each pixel include the predicted probability for each cell, 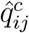, where *c* = 1, …, *C*. Since there are multiple background probabilities, we average them per pixel to form the predicted background probability:

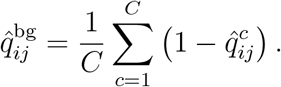

The final segmentation result for a pixel is determined by selecting the highest probability among the background and each cell in the patch:

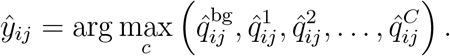

### Training Details

#### Training Objective for the Foreground Predict Net

The training objective for the Foreground Predict Net begins with the definition of foreground and background samples. We define 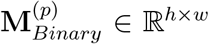 to be the binary mask for all nucleus in the patch. For the foreground, pixels located within the nuclei regions 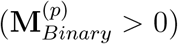 are utilized as ground truth positive samples. This ensures that the model accurately identifies areas that are indeed part of the cells.

To define the negative samples, the nuclei segmentation mask is manipulated to generate 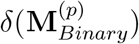, where δ represents a dilation operator. This operator dilates the nuclei masks using a 10*×*10 kernel over four iterations by default. Subsequently, the average expression on the zero-area of the dilation mask is calculated using the formula:

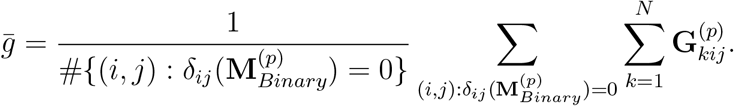

The background is then defined as **B**^(*p*)^ :

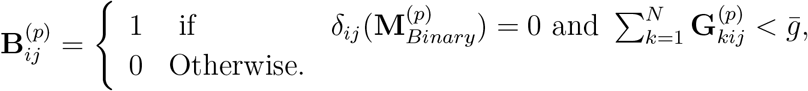

This formulation ensures that regions without significant transcriptional activity are correctly identified as non-cell areas. Pixels that do not fall within the nuclei or background regions are excluded from the training process. Recall that 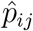 is the predicted probability that pixel (*i, j*) belongs to the foreground. The weighted Cross-Entropy (CE) loss for training the Foreground Predict Net *L*_*fg*_ on a single patch is then formulated as:

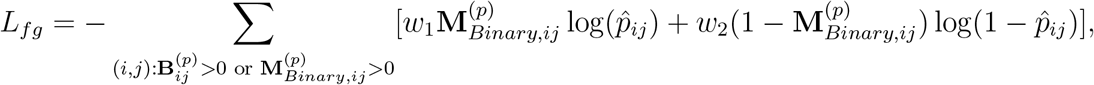

where *w*_1_ and *w*_2_ are the weighted parameters of the foreground region and background region, respectively. This selective training approach enhances the model’s ability to distinguish between cellular and non-cellular regions.

#### Training Objective for the Cell Predict Net

The training for the Cell Predict Net relies on the predictions of the Foreground Predict Net. Similar to the training process above, for each cell we designate the nuclei region as the positive sample, while the background predicted by the Foreground Predict Net 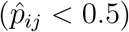 is treated as the negative sample. The other region will be dropped at the training stage. Suppose *C* represents the number of nuclei in the nuclei segmentation mask **M**^(*p*)^ and 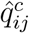 denotes the predicted segmentation probability for cell *c* in the patch, The final weighted CE loss for training the Cell Predict Net, *L*_*cell*_ on a single patch is then formulated as follows:

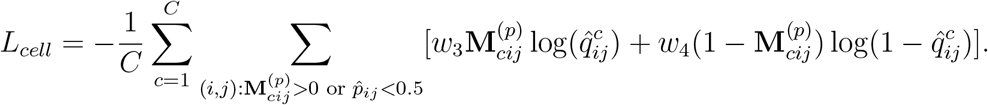

In this formulation, **M**_*p,c*_(*i, j*) is the binary mask indicating whether the spot (*i, j*) belongs to the nuclei of cell *c*. The weights *w*_3_ and *w*_4_ are balancing factors that adjust the contribution of positive and negative samples to the loss function. This weighted CE loss ensures that the model learns to accurately distinguish between cell interior and predicted background regions, thus refining the segmentation boundaries.

#### Other details

An essential aspect of our training process is the influence of the dilation parameters for defining the background area. Adjusting this parameter allows us to balance cell size and the number of included transcripts per cell, ensuring that our model can adapt to various tissue types and experimental conditions. The training process is expedited by the model’s relatively simple architecture and the limited number of parameters, allowing for rapid convergence. For optimization, we employ the Adam optimizer with a learning rate of 10^−4^, *β*_1_ = 0.9, *β*_2_ = 0.999, and a weight decay of 10^−4^ across different datasets. The weighted parameters *w*_1_, *w*_2_, *w*_3_, *w*_4_ are generally set to 1. For the Stereo-seq dataset, *w*_1_ and *w*_3_ are set to 5 as their default values. The postprocessing procedure for UCS adheres closely to the strategies employed by BIDCell that combine with several morphological operations.

### Scaled softmask and integrating the marker gene information

To enhance the segmentation of elongated cells without compromising the accuracy of normal cells, we propose a refined approach to scaling the softmask. This process begins with fitting a minimal enclosing ellipse around the nuclei of each cell. The major and minor axes of the ellipse are denoted as *a* and *b*, respectively. The eccentricity *e* of the ellipse is then calculated using the formula

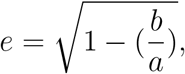

where *a* and *b* represent the lengths of the major and minor axes. Following this, the distance between each pixel and the closest point on the nucleus is decomposed into two components along the major and minor axes, denoted as *d*_*a*_ and *d*_*b*_. These distances are then scaled by the square root of the eccentricity. Specifically, the distance along the major axis *d*_*a*_ is multiplied by 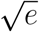, and the distance along the minor *d*_*b*_ is divided by 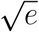. This recalibration results in the weight of each pixel along the major axis becoming larger and along the minor axis becoming smaller (Supplementary Fig. 3). If the nucleus is nearly circular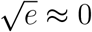, the distances remain largely unchanged, thus preserving the segmentation accuracy for normal cells.

In addition to adjusting the softmask, we integrate marker gene information into the model. This integration starts with annotating the nuclei to obtain cell types. For each cell, a mask is generated where each pixel is assigned a value based on the presence of marker genes in the transcript map: if a marker gene is present at a given pixel, it is assigned a value of 1; otherwise, it is assigned a value of 0. This marker gene mask is then used as an additional input to the Cell Predict Net. An additional cross-entropy loss function is introduced. This loss function penalizes discrepancies between the predicted segmentation and the marker gene mask, thereby guiding the model to incorporate the marker gene data into its segmentation predictions. Note that scaling the softmask for elongated cells and integrating marker gene information are designed to complement the normal segmentation procedure without significantly altering it.

### Addressing Gene Variability and Tuning the hyperparameters

For datasets encompassing the whole transcriptome, such as Stereo-seq and NanoString CosMx, which include more than 10,000 genes, we identify the top 1,000 highly variable genes (HVGs) after filtering out control genes. This step is crucial for focusing on the most informative genes and reducing computational complexity. Given that the Stereo-seq Mouse Brain dataset captures an entire adult mouse brain slice (an area of approximately 5.3 × 7.0 *mm*^2^) with a resolution of 0.5 *µ*m, the transcript map is significantly larger compared to datasets such as 10X Xenium and Vizgen MERSCOPE.

It is essential to note that different regions of the Stereo-seq dataset exhibit distinct sets of HVGs due to the diversity in cell types (Supplementary Fig. 4). This observation underscores a potential flaw in previous transcriptomics methodologies that apply the same set of HVGs across the entire tissue, potentially overlooking regional variations in gene expression. To address this, we segment the Stereo-seq dataset into small datasets of 1200 × 1200 pixels and train a separate model on each small dataset. Similarly, for the NanoString CosMx dataset, we train models on each field of view (FOV) to accommodate the variability in gene expression across different regions of the tissue.

The UCS workflow is detailed in the supplementary materials (Supplementary Fig. 5). Regarding hyperparameters, we primarily adjust the number of training epochs based on the size of the SST dataset and the parameters for the dilation kernel and iterations, which influence the perceived cell size.

### Downstream analysis

For the downstream analysis of our SST data, we employed several specialized software tools. We used Single-cell Variational Inference (SCVI) [21] [22] (https://scvi-tools.org/) to transfer labels from the scRNA-seq dataset to the cells in the SST dataset based on segmentation. For cellular level downstream analysis, we utilized Scanpy [23] (https://scanpy.readthedocs.io/en/stable/) and Squidpy [24] (https://squidpy.readthedocs.io/en/stable/), which are popular tools for single-cell gene expression analysis and spatial pattern detection, respectively. Additionally, we used Bento [17] (https://github.com/ckmah/bento-tools) for subcellular gene classification. These tools collectively enhance our analysis and interpretation of high-resolution spatial gene expression data.

### Parameter setting for method comparison

We used several methods to benchmark against our proposed method. We employed SCS on Stereo-seq using the default parameters. For BIDCell, due to its requirement for extensive additional data, such as negative and positive marker genes, we limited its application to the Xenium Breast Cancer dataset. We followed the official instructions (https://www.10xgenomics.com/jp/analysis-guides/using-baysor-to-perform-xenium-cell-segmentation) detailed in the 10x Genomics analysis guide to run Baysor. Cellpose was used for segmenting the nuclei staining using the “cyto” model. Last, we generated Voronoi diagrams using the py-clesperanto Python package (https://github.com/clEsperanto/pyclesperanto_prototype).

## Data availability

All datasets used in this work are publicly available through online sources.

- Xenium Human Breast Cancer and the scRNA Reference using the adjacent tissues (https://www.10xgenomics.com/products/xenium-in-situ/preview-dataset-human-breast)
- Xenium Mouse Brain (https://www.10xgenomics.com/datasets/fresh-frozen-mouse-brain-replicates-1-standard)
- Xenium Human Lung Cancer (https://www.10xgenomics.com/datasets/preview-data-ffpe-human-lung-cancer-with-xenium-multimodal-cell-segmentation-1-standard)
- Stereo-seq Mouse Brain (https://db.cngb.org/stomics/mosta/download/)
- NanoString CosMx Human Pancreas (https://nanostring.com/products/cosmx-spatial-molecular-imager/ffpe-dataset/cosmx-smi-human-pancreas-ffpe-dataset/)
- Vizgen MERFISH Mouse Brain S2R1 (https://info.vizgen.com/mouse-brain-map)
- Vizgen MERSCOPE Ovarian Cancer 3 (https://info.vizgen.com/ffpe-showcase)

## Code availability

The UCS software is available at https://github.com/YangLabHKUST/UCS.

## Acknowledgements

We acknowledge the following grants: Hong Kong Research Grant Council grants nos. 16301419, 16308120, 16307221, 16307322, and 16302823, The Hong Kong University of Science and Technology Startup Grants R9405 and Z0428 from the Big Data Institute.

## Notes

### Competing Interest Statement

The authors have declared no competing interest.

