## Supplementary for "UCS: a unified approach to cell segmentation for subcellular spatial transcriptomics"

### Supplementary Figures

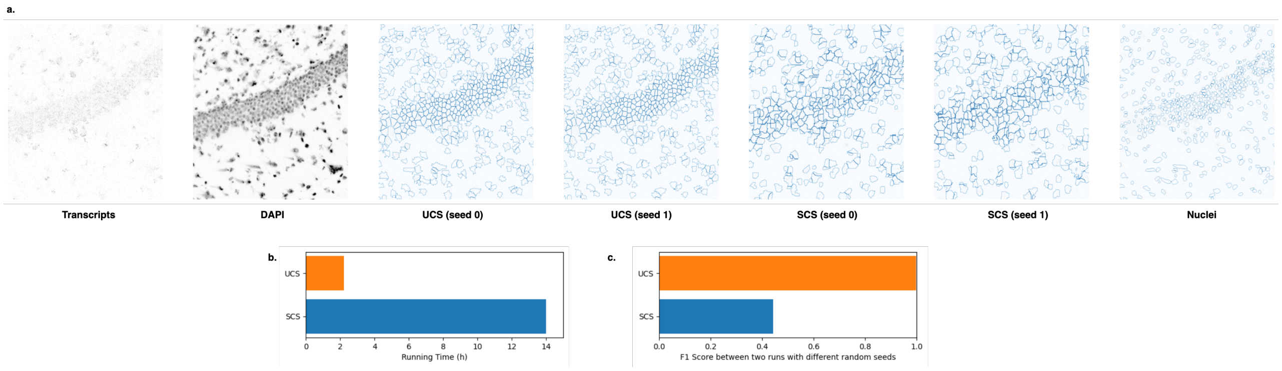


Figure 1: Comparison of UCS and SCS training on a 1200x1200 patch of the Stereo-seq dataset. **a**. Visualization of segmentation results for UCS and SCS using different random seeds. **b**. Comparison of running times for both methods. **c**. F1 score comparison between segmentation results from two different runs.

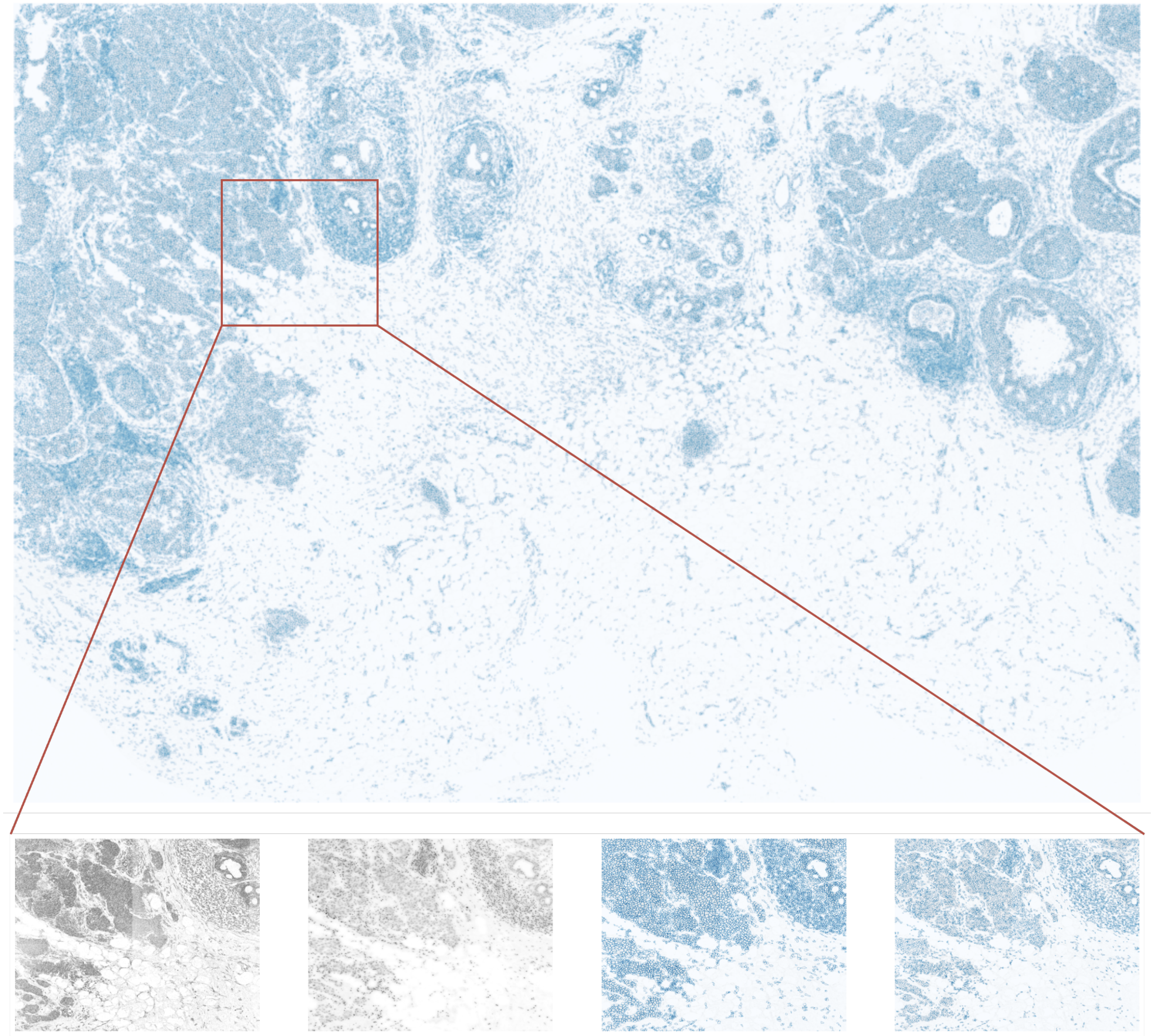


Figure 2: Evaluation of UCS on Xenium Breast Cancer, Replicate 2. The model was trained exclusively on Xenium Breast Cancer, Replicate 1, to demonstrate its robustness.

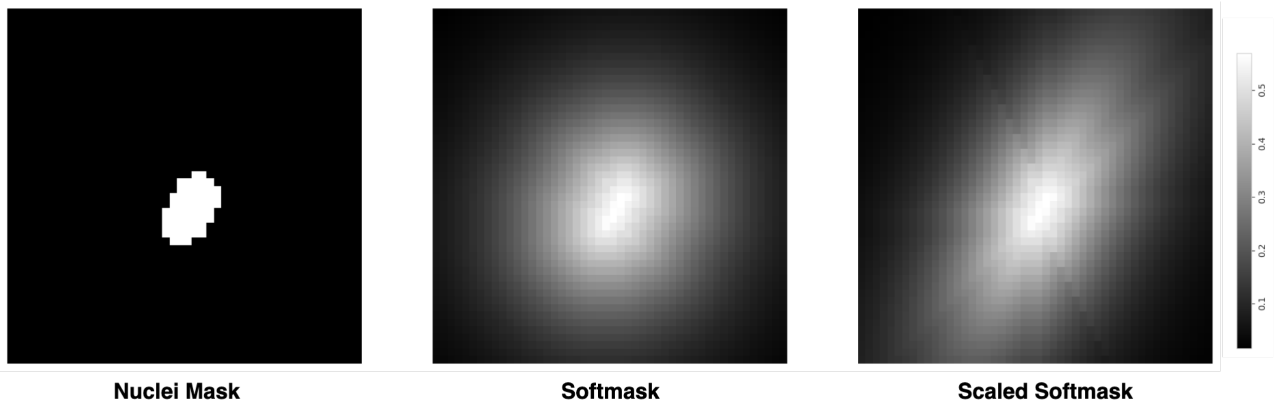


Figure 3: Softmask used by UCS. Left: Raw nuclei mask. Middle: Softmask. Right: Scaled softmask preserving nuclear morphology

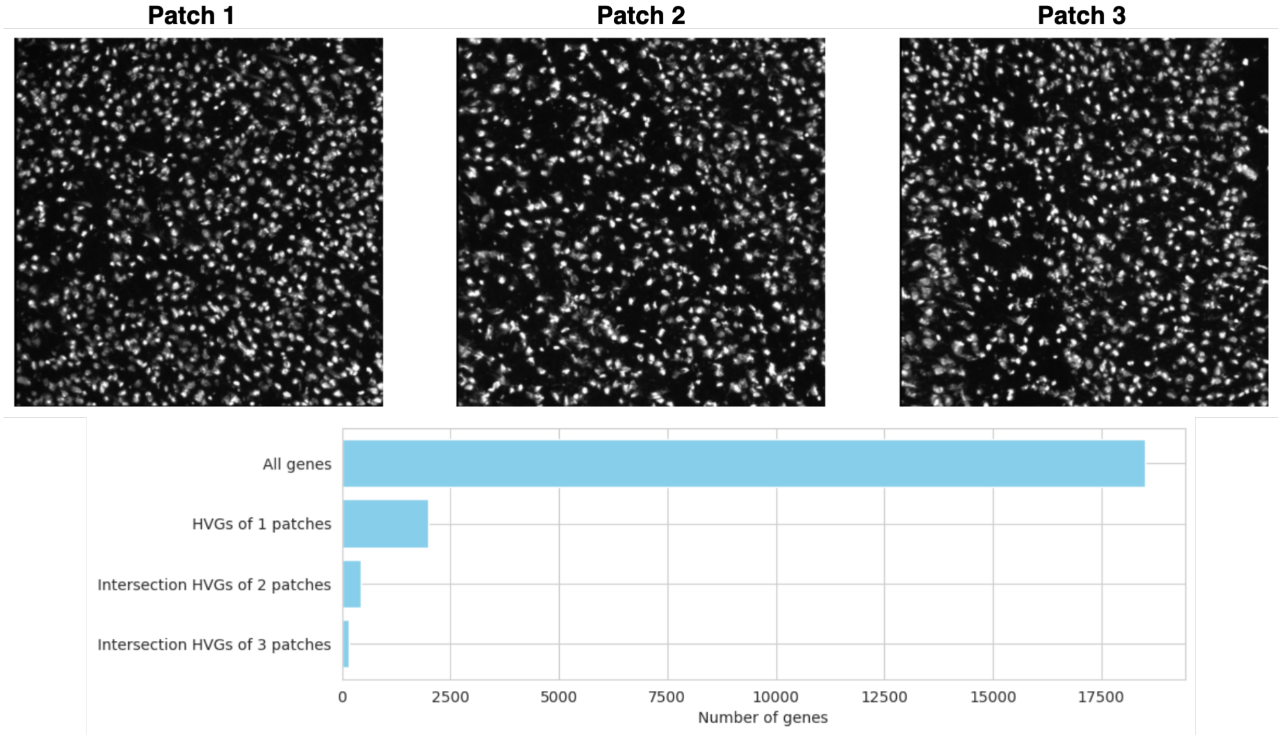


Figure 4: Highly Variable Genes (HVGs) Vary Across Different Tissue Patches. We selected three patches from the Stereo-seq dataset, each containing a sufficient number of cells and located in distinct regions of the tissue. For each patch, 2000 HVGs were identified. The overlap among the HVGs from the three patches reveals that only 161 genes are common, highlighting the variability in gene expression across different tissue regions.

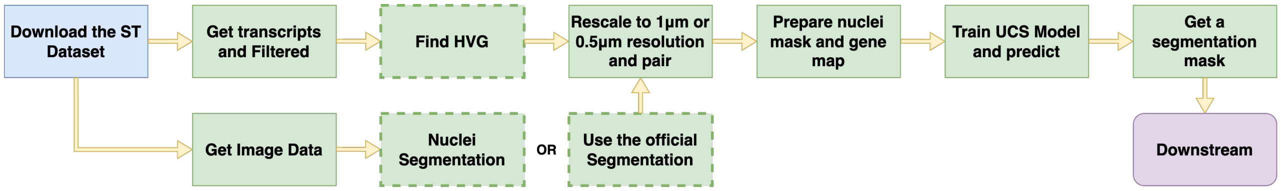


Figure 5: Workflow of UCS

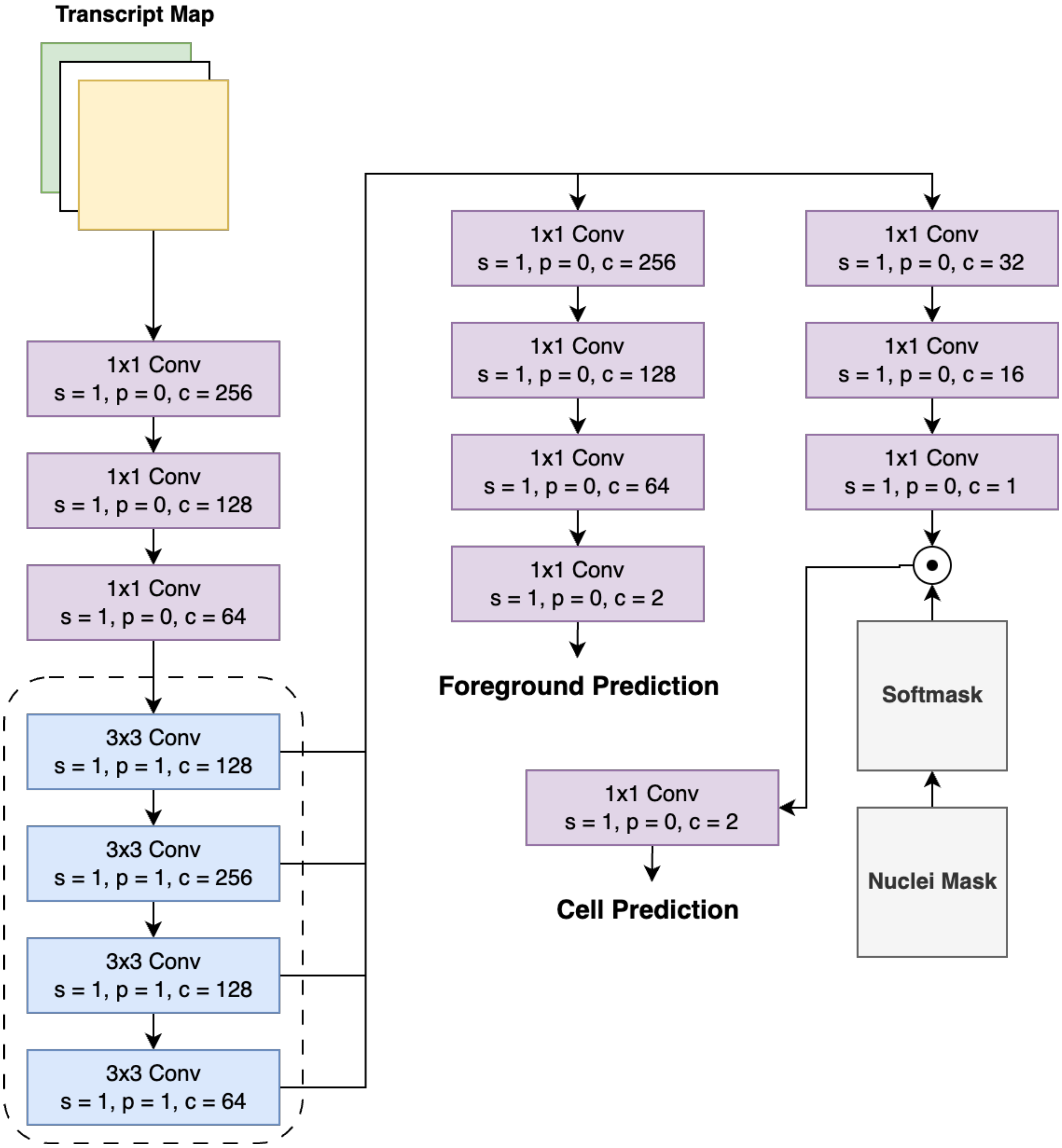


Figure 6: Architecture of the UCS Network
